# SINTAX: a simple non-Bayesian taxonomy classifier for 16S and ITS sequences

**DOI:** 10.1101/074161

**Authors:** Robert C. Edgar

## Abstract

Metagenomics experiments often characterize microbial communities by sequencing the ribosomal 16S and ITS regions. Taxonomy prediction is a fundamental step in such studies. The SINTAX algorithm predicts taxonomy by using *k*-mer similarity to identify the top hit in a reference database and provides bootstrap confidence for all ranks in the prediction. SINTAX achieves comparable or better accuracy to the RDP Naive Bayesian Classifier with a simpler algorithm that does not require training. Most tested methods are shown to have high rates of over-classification errors where novel taxa are incorrectly predicted to have known names.

## Introduction

Sequencing of tags such as the ribosomal 16S gene and fungal internal transcribed space (ITS) region is a popular method for surveying microbial communities. Recent examples include the Human Microbiome Project (HMP Consortium, 2012) and a survey of the *Arabidopsis* root microbiome (Lundberg *et al.*, 2012). A fundamental step in such studies is to predict the taxonomy of sequences found in the reads. The most popular method is currently the RDP Naive Bayesian Classifier (Wang *et al.*, 2007) (hereafter RDP). Additional taxonomy prediction methods are supported by QIIME (Caporaso *et al.*, 2010) and mothur (Schloss *et al.*, 2009).

### Reference databases

Taxonomy prediction requires a reference database containing sequences with taxonomy annotations. Authoritative prokaryotic sequence classifications exist for at most the ~12,000 named species belonging to ~2,300 genera which represent only a tiny fraction of extant species (Yarza *et al.*, 2014). Available databases include the RDP training sets, the full RDP database (RDPDB) (Maidak *et al.*, 2001), SILVA (Pruesse *et al.*, 2007), Greengenes (DeSantis *et al.*, 2006) and UNITE (Kõljalg *et al.*, 2013). The RDP 16S training set v16 (RTS) has 13,212 sequences belonging to 2,126 genera while the RDP Warcup ITS training set (Deshpande *et al.*, 2015) v2 has 18,878 sequences belonging to 8,551 species. The RDP training sets contain only sequences with authoritative names and are therefore much smaller than SILVA, Greengenes and UNITE which include environmental sequences. SILVA v123 has 1.8M small subunit ribosomal RNA sequences; v114 was estimated to contain ~94,000 genera (Yarza *et al.*, 2014). Greengenes v13.5 has 1.8M 16S sequences. UNITE release 01.08.2015 has 476k ITS sequences representing ~71,000 species. Most taxonomy annotations in SILVA and Greengenes are predictions obtained by computational and manual analyses which are primarily based on trees predicted from multiple alignments (McDonald *et al.*, 2012; Yilmaz *et al.*, 2014); in RDPDB most annotations are predicted by RDP. In the 16S databases (RDPDB, SILVA and Greengenes), no attempt is made to classify unnamed groups, while UNITE assigns numerical “species hypothesis” identifiers to unnamed clusters. By default, QIIME uses a subset of Greengenes clustered at 97% identity (GGQ, containing 99k sequences in v13.8), and mothur recommends a subset of SILVA (SILVAM, containing 172k sequences in v123). The RDP web site and stand-alone software use the RDP training sets.

### Database coverage and novel taxa

If a query sequence is found in the database, its taxonomy is naively given by the reference annotation. This prediction may wrong if the database has annotation errors or multiple species are identical over the sequenced region, which often happens with short tags such as the popular V4 hypervariable region of 16S. The latter scenario cannot be reliably identified by checking the database for other identical sequences because the reference data may be incomplete. If the query sequence is not found in the database then prediction is more difficult. For example, using a 95% identity threshold for clustering full-length 16S sequences was found to give groups that best approximate genera (Yarza *et al.*, 2014). Thus, if a 16S sequence has 95% identity with a database hit, it might be in the same genus but since identity correlates only approximately with taxonomic rank it could belong only to the same family or same class. Or, it could belong to the same species if there is atypically large variations between paralogs or strains. From this perspective, the task of taxonomy prediction is to estimate the *lowest common rank* (LCR) between the query and the database. A query rank *r* is *known* if *r* ≥ LCR, i.e. at least one member of its clade is present in the reference database (regardless of whether it is named) and *novel* if *r* < LCR. The *coverage* of a reference database at a given rank with respect to a set of query sequences is the fraction of queries that are known and *novelty* = (1 – coverage) is the fraction of queries that are novel. The mean top-hit identity (MTI) between query sequences and their top hits can be used as an approximate indication of coverage. To obtain typical query sets, I constructed OTUs at 97% identity using UPARSE (Edgar, 2013) from V4 reads of human gut, mouse gut and soil communities respectively (Kozich *et al.*, 2013) and ITS reads of a soil fungal community (Schmidt *et al.*, 2013). MTIs of these samples vs. commonly-used reference databases are shown in Table 1. All V4 samples have MTI<95% with RTS, suggesting that many, perhaps most, OTUs belong to novel genera, especially in soil (MTI=88%).

**Table 1.**
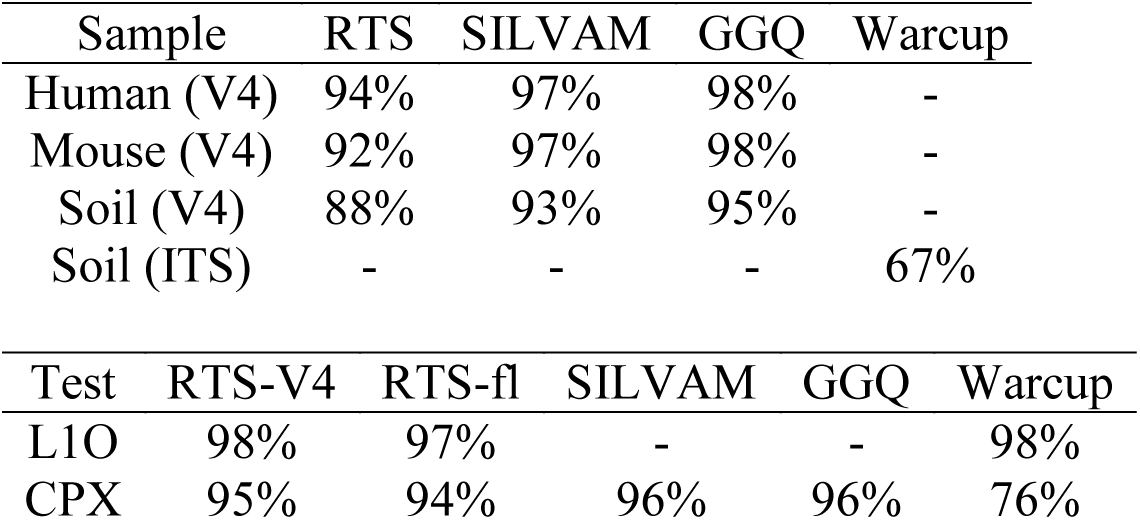
Mean top-hit identities

| Sample | RTS | SILVAM | GGQ | Warcup |
| --- | --- | --- | --- | --- |
| Human (V4) | 94% | 97% | 98% | - |
| Mouse (V4) | 92% | 97% | 98% | - |
| Soil (V4) | 88% | 93% | 95% | - |
| Soil (ITS) | - | - | - | 67% |

| Test | RTS-V4 | RTS-fl | SILVAM | GGQ | Warcup |
| --- | --- | --- | --- | --- | --- |
| L1O | 98% | 97% | - | - | 98% |
| CPX | 95% | 94% | 96% | 96% | 76% |

### RDP leave-one-out validation

RDP was tested on 16S and ITS sequences using leave-one-out validation (Wang *et al.*, 2007; Deshpande *et al.*, 2015) where one query sequence is extracted from the training set (RTS and Warcup, respectively) and classified using the remaining sequences as a reference. Accuracy (*Acc*_RDP_) is calculated as the fraction of sequences that are correctly classified. Roughly half (1,119 / 2,472) of the genera in RTS are singletons, i.e. have exactly one training sequence, while about a quarter (2,258 / 8,548) of the species in Warcup are singletons, comprising 8% (16S) and 13% (ITS) of the training sequences. A singleton cannot be classified correctly in a leave-one-out test because no training sequences are left for its clade so that the maximum achievable *Acc*_RDP_ by an ideal algorithm is the fraction of non-singleton taxa, i.e. 92% for 16S genus and 87% for ITS species, rather than 100% as would usually be expected for an accuracy measure. The average number of non-singleton training sequences is 9 per genus in RTS and 14 per species in Warcup which suggests that correct classification should be relatively easy for most queries, while in practice many genera will be novel, and taxa that are rare in the database may be common in the query set and vice versa. Also, all predictions are included in *Acc*_RDP_ regardless of their bootstrap confidence values rather than using the authors’ recommended parameters (here, 80% cutoff) as would usually be expected for a benchmark test. In summary, the RDP leave-one-out test does not model typical query datasets and *Acc*_RDP_ does not give a realistic estimate of accuracy by any conventional definition.

## Methods

### Performance metrics

Sensitivity should be measured as the fraction of *known* queries that are correctly identified so that the highest achievable sensitivity by an ideal algorithm is 100%. If novel queries were also counted then sensitivity <100% would reflect an opaque combination of low database coverage and failures to correctly predict known taxa, as with *Acc*_RDP_. It is useful to distinguish two types of false positive error: *misclassifications*, where an incorrect name is predicted for a known rank, and *over-classifications*, where a name is predicted for a novel rank. For a given query set, reference database and taxonomic rank let *N*_*known*_ and *N*_*novel*_ be the number of queries with known and novel taxa respectively. Let *TP* be the number of correct predictions, *FP*_*mis*_ be the number of misclassification errors and *FP*_*over*_ be the number of over-classification errors. The total number of queries is *N* = *N*_*known*_ + *N*_*novel*_. The following accuracy metrics can now be defined:

Sensitivity = TP / *N*_*known*_,

Misclassification rate = MC = *FP*_*mis*_ / *N*_*known*_,

Over-classification rate = OC = *FP*_*over*_ / *N*_*novel*_,

Errors per query = EPQ = (*FP*_*mis*_ + *FP*_*over*_) */ N*.

To a first approximation, we might expect misclassification and over-classification rates to be similar on different datasets because these measures reflect intrinsic characteristics of an algorithm independent of the data while EPQ, the measure that is typically of most interest in practice, will strongly depend on database coverage (equivalently, on query novelty). For example, if a query set contains mostly known sequences, we would expect errors to be rare and dominated by misclassifications, while if a query set is highly novel then there may be many overclassifications. If these expectations are correct, then values of MC and OC measured on a benchmark test will be similar to those obtained on biological data in practice while EPQ will be similar only if the benchmark has similar rates of novel taxa.

### Clade partition cross-validation (CPX)

If high ranks are usually known but low ranks are often novel, then a benchmark test should contain a mix of known and novel taxa at low ranks so that both MC and OC can be measured. This can be achieved by *clade partition cross-validation* (CPX), as follows. Clades at a given rank *r*_*part*_ from a reference database are partitioned so that a randomly-chosen half of the daughter groups in a given clade are assigned to the query set and the other half to the reference set so that ranks below *r*_*part*_ are always novel. For example, if *r*_*part*_ = family then half of the genera for a given family are assigned to the query and half to the reference set. Singletons are always assigned to the query set, so are always novel while non-singletons are always known. For this work, I used *r*_*part*_ = family and *r*_*part*_ = genus and calculated performance metrics from the combined predictions on both query-reference pairs.

### SINTAX algorithm

For a query sequence *Q* and reference database *R* the SImple Non-Bayesian TAXonomy (SINTAX) algorithm proceeds as follows. Let *W*(*Q*) be the set of *k*-mers in *Q* where *k* = 8 by default. In one iteration, a random sub-sample *w*_*s*_(*Q*) of size *s* is extracted from *W*(*Q*) where *s* = 32 by default. Sub-sampling is performed with replacement. For each reference sequence *r* ∈ *R*, the number of words in common is *U*^*subset*^(*r*) = |*w*_*s*_(*Q*) ⋂ *W*(*r*)|. The top hit *T* by *k*-mer similarity is identified as *T* = *argmax*(*r*) *U*^*subset*^(*r*) and the taxonomy is taken from the annotation of *T*. By default, 100 iterations are performed. For each rank, the name that occurs most often is identified and its frequency is reported as its bootstrap confidence. SINTAX is similar to the RDP algorithm except (a) the taxonomy in each iteration is identified from the top *k*-mer hit for the sub-sample rather than the most probable taxonomy according to the naive Bayesian calculation and (b) a fixed-size subset of 32 *k*-mers is used for bootstrapping. RDP uses a subset of size |*Q*|/*k*, the number of non-overlapping *k*-mers, because overlapping *k*-mers are not independent. SINTAX uses a fixed-size subset to compensate for a problem that arises with longer sequences. For a given query sequence, consider reference sequences ranked using all *k*-mers, i.e. in order of decreasing *U*^*all*^(*r*) = |*W*(*Q*) ⋂ *W*(*r*)|. This gives a list of taxonomies sorted by decreasing *U*^*all*^. Let *C*^*all*^_1_ be the top taxonomy and *C*^*all*^_2_ the second-ranked taxonomy with similarities *U*^*all*^_1_ and *U*^*all*^_2_ respectively using all *k*-mers, and similarities *U*^*subset*^_1_ and *U*^*subset*^_2_ using the subset in a given iteration. If *U*^*all*^_1_ ≫ *U*^*all*^_2_ then *U*^*subset*^_1_ will be greater than *U*^*subset*^_2_ in most or all iterations and *C*_1_ will therefore have high bootstrap confidence. Conversely, if *U*^*all*^_1_ is only slightly greater than *U*^*all*^_2_, then it is more likely that the order of the top two taxonomies will be reversed in *U*^*subset*^ order and the bootstrap confidence of *C*_1_ will then be lower. The bootstrap confidence thus correlates with the difference *U*^*all*^_1_ – *U*^*all*^_2_, giving an indication of how much closer the top taxonomy is to the query than the second-ranked taxonomy. As the sequence length |*Q*| increases, the number of non-overlapping *k*-mers |*Q*|/*k* increases. Using a larger subset reduces fluctuations under sub-sampling so that the taxonomy order defined by *U*^*subset*^ converges on the order defined by *U*^*all*^, and in particular *C*_1_ is the top-ranked taxonomy more often. The bootstrap confidence of *C*_1_ therefore tends to increase for longer sequences, regardless of whether it is correct. In other words, as the sequence length increases, using a subset of size |*Q*|/*k* tends to give high bootstrap confidence to the top *k*-mer hit for any query sequence. When the query is novel, *C*_1_ is always wrong and the over-classification rate therefore increases with longer sequences. This problem is mitigated by using a fixed subset size of 32. See Supp. Table 1 for a comparison of SINTAX with *s*=32 and *s*=1/*k* vs. RDP.

## Results

I tested SINTAX v1.0, standalone RDP v2.12, QIIME v1.9.1 and mothur v1.36.1. The QIIME *assign_taxonomy.py* script was run with options *-m uclust* (*Quc*, the default), -*m sortme* (*Qsm*),-*m blast* (*Qblast*) and -*m rdp* (*Qrdp*, a wrapper for standalone RDP that sets a bootstrap cutoff of 50% by default). The mothur *classify.seqs* command was run with *method=wang* (*Mrdp*, the default, a re-implementation of the RDP algorithm) and *method=knn* (*Mknn*).

### Leave-one-out validation

Results for leave-one-out testing of SINTAX and RDP are shown in Table 2 and Supp. Table 1. The *Acc*_RDP_ metric is shown together with Sensitivity, OC and MC for genus (16S) or species (ITS). At phylum rank, EPQ is given as the measure of error rate since almost all phyla are known so OC cannot be measured reliably and MC ≈ EPQ.

**Table 2.** Leave-one-out test results

| $r(b)$ | metric | RTS (V4) | | RTS (fl) | | Warcup | |
| --- | --- | --- | --- | --- | --- | --- | --- |
|  |  | RDP | SINTAX | RDP | SINTAX | RDP | SINTAX |
| p(0) | $Acc_{RDP}$ | 99.7 | 99.7 | 99.5 | 99.8 | 99.9 | 100.0 |
| $t(0)$ | $Acc_{RDP}$ | 80.4 | 80.5 | 85.6 | 85.6 | 73.9 | 75.6 |
| p(80) | Sens. | 99.3 | 99.5 | 99.7 | 99.7 | 99.9 | 100.0 |
| p(80) | EPQ | 0.3 | 0.0 | 0.5 | 0.0 | 0.0 | 0.0 |
| $t(80)$ | Sens. | 78.9 | 77.1 | 92.1 | 81.1 | 79.8 | 75.2 |
| $t(80)$ | OC | 28.5 | 28.3 | 48.0 | 16.8 | 42.2 | 25.6 |
| $t(80)$ | MC | 3.5 | 3.6 | 2.8 | 1.2 | 7.9 | 3.2 |
| $t(80)$ | EPQ | 5.9 | 5.9 | 7.1 | 2.7 | 12.7 | 6.4 |

### Clade partition cross-validation

Results for CPX testing are shown in Table 3 and Supp. Table 2. SINTAX is compared to other algorithms at genus and phylum ranks using the default database for each method. Against RDP, a bootstrap cutoff of 80% is used as this is the value recommended by the RDP authors. SINTAX and RDP are observed to have similar performance on V4 while SINTAX has substantially lower error rates on full-length 16S and ITS due to its lower over-classification rates. The ITS phylum sensitivity of SINTAX (98.3%) is notably better than RDP (81.8%).

**Table 3.** CPX test results

| Algo | Cutoff | OC<sub>g</sub> | MC<sub>g</sub> | Sens<sub>g</sub> | EPQ<sub>g</sub> | Sens<sub>p</sub> | EPQ<sub>p</sub> |
| --- | --- | --- | --- | --- | --- | --- | --- |
| SINTAX | 80 | 23.1 | 2.7 | 82.2 | 14.4 | 96.9 | 0.1 |
| RDP | 80 | 21.3 | 3.0 | 82.1 | 13.5 | 90.0 | 0.2 |

**RTS (full-length)**
| Algo | Cutoff | OC<sub>g</sub> | MC<sub>g</sub> | Sens<sub>g</sub> | EPQ<sub>g</sub> | Sens<sub>p</sub> | EPQ<sub>p</sub> |
| --- | --- | --- | --- | --- | --- | --- | --- |
| SINTAX | 80 | 12.2 | 0.4 | 84.7 | 6.8 | 98.0 | 0.0 |
| RDP | 80 | 40.3 | 1.6 | 94.0 | 22.5 | 92.7 | 0.9 |

**Warcup (ITS)**
| Algo | Cutoff | OC<sub>g</sub> | MC<sub>g</sub> | Sens<sub>g</sub> | EPQ<sub>g</sub> | Sens<sub>p</sub> | EPQ<sub>p</sub> |
| --- | --- | --- | --- | --- | --- | --- | --- |
| SINTAX | 80 | 14.4 | 0.8 | 95.6 | 6.4 | 98.3 | 0.3 |
| RDP | 80 | 67.8 | 1.0 | 91.2 | 41.3 | 81.8 | 10.8 |

**GGQ (V4)**
| Algo | Cutoff | OC<sub>g</sub> | MC<sub>g</sub> | Sens<sub>g</sub> | EPQ<sub>g</sub> | Sens<sub>p</sub> | EPQ<sub>p</sub> |
| --- | --- | --- | --- | --- | --- | --- | --- |
| SINTAX | 80 | 28.0 | 5.9 | 75.7 | 23.0 | 91.4 | 0.4 |
| SINTAX | 50 | 56.2 | 10.2 | 83.7 | 45.7 | 95.7 | 2.0 |
| *Quc* | - | 48.4 | 9.7 | 77.3 | 39.6 | 72.8 | 0.4 |
| *Qsm* | - | 45.7 | 7.7 | 76.1 | 37.1 | 73.3 | 0.3 |
| *Qblast* | - | 87.4 | 17.1 | 82.7 | 71.4 | 90.2 | 8.4 |
| *Qrdp* | 50 | 49.4 | 9.6 | 81.7 | 40.3 | 95.0 | 1.1 |

**SILVAM (V4)**
| Algo | Cutoff | OC<sub>g</sub> | MC<sub>g</sub> | Sens<sub>g</sub> | EPQ<sub>g</sub> | Sens<sub>p</sub> | EPQ<sub>p</sub> |
| --- | --- | --- | --- | --- | --- | --- | --- |
| SINTAX | 80 | 35.7 | 5.3 | 80.9 | 21.8 | 98.3 | 0.4 |
| *Mrdp* | 80 | 30.8 | 2.4 | 75.8 | 17.8 | 97.1 | 0.1 |
| *Mknn* | - | 22.3 | 0.8 | 61.5 | 12.5 | 98.0 | 0.4 |

### High over-classification rates

A high rate of over-classifications at low rank was found for all methods on all tests ranging from a minimum genus OC of 12.2% (SINTAX on RTS V4 with 80% bootstrap cutoff) to a maximum of 87% (*Qblast* on GGQ V4). The QIIME OC rates were especially high. The lowest OC of any QIIME method (tested on V4 with its default GGQ reference database) was 48.4% (*Qsm*). A high OC rate is of practical importance if a significant number of novel genera are present in the query set. Table 1 shows that the OTUs for the V4 soil data have mean identity 95% or less with all of the default reference databases, implying that half or more probably have novel genera. On a query set with 50% novel genera, an algorithm will have an overall false-positive (FP) rate of 0.5×OC + 0.5×MC. At genus rank, using OC and MC values measured on the CPX test, this gives a FP rate of 13% for RDP and SINTAX at 80% bootstrap using RTS, 29% for the default QIIME method *Quc* and 52% for *Qblast*.

## Discussion

SINTAX achieves comparable (V4) or better (full-length 16S and ITS) accuracy to RDP. SINTAX is conceptually simple: it finds the top *k*-mer hit, and the evidence supporting a prediction can be presented as a list of reference taxonomies with their *k*-mer similarities to the query sequence. The RDP algorithm is more opaque, making it difficult to review the evidence supporting a prediction. The posterior probabilities calculated according to the naive Bayesian theory are astronomically small for correct predictions, typically in the range 10–18 to 10–24. Ideally, a posterior would be an estimate of the probability that a taxonomy is correct and would be ~1 for a correct prediction, but here the probabilities are wrong by twenty or so orders of magnitude, necessitating post-hoc bootstrapping to obtain a useable confidence measure. SINTAX and RDP have similar accuracy when both use sub-sample size |*Q*|/*k* for bootstrapping (Supp. Table 2) in which case the algorithms are essentially the same except for the score for sorting taxonomies. This suggests that the naive Bayesian approach could be interpreted as an approximation to finding the top *k*-mer hit.

The high measured over-classification rates indicate an unsolved problem with novel taxa, which is readily explained by sparse reference data (Fig. 1). At high ranks, this may not be important because novel phyla are rare, but at low ranks novel taxa are common, especially in communities that are less studied or difficult to culture (e.g. extremophiles) or highly diverse (e.g. soil).

**Figure 1.**
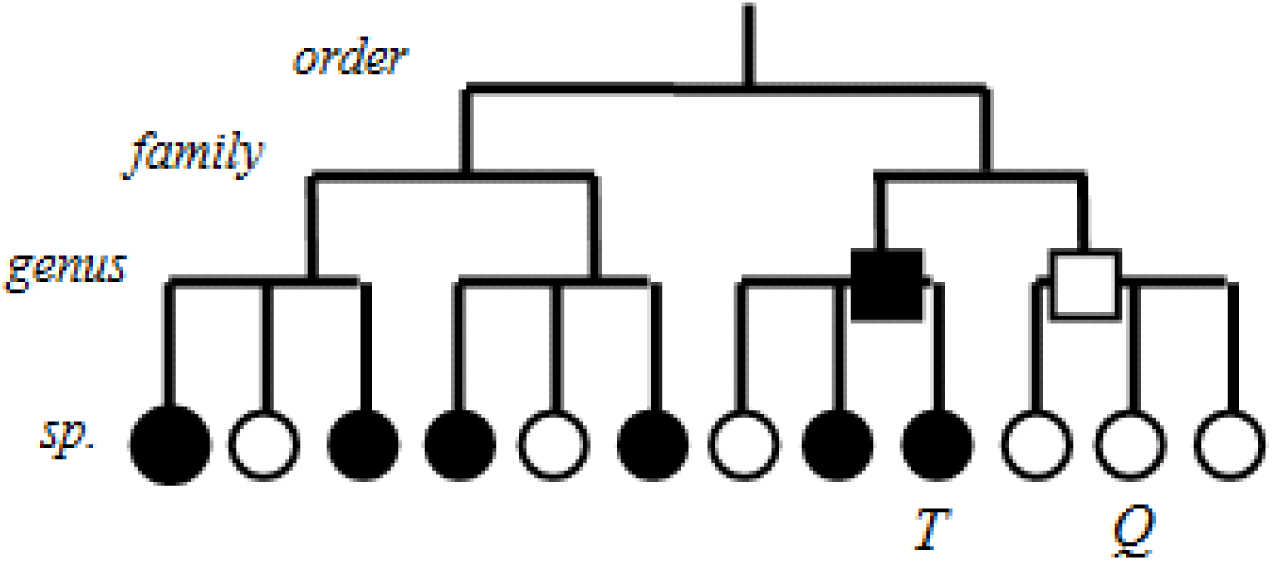
A sparse reference database induces over-classification errors. *Q* is the query and *T* is the top hit. Black circles denote known species, white circles novel species. If *Q* belongs to a novel genus (white square), the top few hits will tend to belong to the most similar known genus (black square). Here, SINTAX will tend to give a high bootstrap confidence to the known genus. Other algorithms which explicitly (or effectively) take a consensus of taxonomies of the most similar reference sequences, such as RDP, will similarly tend to make over-classification errors.

On full-length 16S sequences, RDP has a measured overclassification rate of 40% (CPX) and 48% (leave-one-out) at the recommended 80% bootstrap cutoff. This result suggests that many of the genus annotations in RDPDB, most of which were predicted by RDP at 80% bootstrap, may be false positives as 47% of the 3.2M RDPDB sequences have top-hit identity <95% with RDPTS, implying that roughly half belong to novel genera. Assuming 47% novel genera and the lower OC value gives an estimate of 0.47×0.40×3.2M = 600k over-classified genera.

---

**Supplementary Table 1.** Leave-one-out test results.

V4 SINTAX cutoff=50
| Rank | AccRDP | OC | MC | Sens | EPQ |
| --- | --- | --- | --- | --- | --- |
| --- | --- | --- | --- | --- | --- |
| p | 99.7% | 50.0% | 0.1% | 99.7% | 0.1% |
| g | 80.3% | 56.4% | 8.2% | 86.6% | 12.7% |

V4 RDP cutoff=50
| Rank | AccRDP | OC | MC | Sens | EPQ |
| --- | --- | --- | --- | --- | --- |
| --- | --- | --- | --- | --- | --- |
| p | 99.7% | 50.0% | 0.1% | 99.7% | 0.1% |
| g | 80.4% | 59.0% | 8.2% | 86.6% | 12.9% |

V4 SINTAX cutoff=80
| Rank | AccRDP | OC | MC | Sens | EPQ |
| --- | --- | --- | --- | --- | --- |
| --- | --- | --- | --- | --- | --- |
| p | 99.7% | 0.0% | 0.0% | 99.5% | 0.0% |
| g | 80.3% | 28.3% | 3.6% | 77.1% | 5.9% |

V4 RDP cutoff=80
| Rank | AccRDP | OC | MC | Sens | EPQ |
| --- | --- | --- | --- | --- | --- |
| --- | --- | --- | --- | --- | --- |
| p | 99.7% | 0.0% | 0.0% | 99.3% | 0.0% |
| g | 80.4% | 28.5% | 3.5% | 78.9% | 5.9% |

V4 SINTAX cutoff=90
| Rank | AccRDP | OC | MC | Sens | EPQ |
| --- | --- | --- | --- | --- | --- |
| --- | --- | --- | --- | --- | --- |
| p | 99.7% | 0.0% | 0.0% | 99.3% | 0.0% |
| g | 80.3% | 19.0% | 2.8% | 71.0% | 4.3% |

V4 RDP cutoff=90
| Rank | AccRDP | OC | MC | Sens | EPQ |
| --- | --- | --- | --- | --- | --- |
| --- | --- | --- | --- | --- | --- |
| p | 99.7% | 0.0% | 0.0% | 99.0% | 0.0% |
| g | 80.4% | 19.3% | 2.4% | 73.3% | 4.0% |

V4 SINTAX cutoff=100
| Rank | AccRDP | OC | MC | Sens | EPQ |
| --- | --- | --- | --- | --- | --- |
| --- | --- | --- | --- | --- | --- |
| p | 99.7% | 0.0% | 0.0% | 97.5% | 0.0% |
| g | 80.3% | 6.9% | 1.1% | 44.4% | 1.6% |

V4 RDP cutoff=100
| Rank | AccRDP | OC | MC | Sens | EPQ |
| --- | --- | --- | --- | --- | --- |
| --- | --- | --- | --- | --- | --- |
| p | 99.7% | 0.0% | 0.0% | 95.9% | 0.0% |
| g | 80.4% | 7.9% | 0.9% | 49.1% | 1.5% |

V3-V5 SINTAX cutoff=50
| Rank | AccRDP | OC | MC | Sens | EPQ |
| --- | --- | --- | --- | --- | --- |
| --- | --- | --- | --- | --- | --- |
| p | 99.8% | 100.0% | 0.1% | 99.8% | 0.1% |
| g | 83.2% | 51.5% | 5.2% | 89.6% | 9.6% |

| V3-V5 RDP cutoff=50 |  |  |  |  |  |
| --- | --- | --- | --- | --- | --- |
| Rank | AccRDP | OC | MC | Sens | EPQ |
| ----- |  |  |  |  |  |
| p | 99.8% | 100.0% | 0.1% | 99.8% | 0.1% |
| g | 83.3% | 66.7% | 6.3% | 91.1% | 12.0% |

| V3-V5 SINTAX cutoff=80 |  |  |  |  |  |
| --- | --- | --- | --- | --- | --- |
| Rank | AccRDP | OC | MC | Sens | EPQ |
| ----- |  |  |  |  |  |
| p | 99.8% | 0.0% | 0.0% | 99.6% | 0.0% |
| g | 83.2% | 21.7% | 1.8% | 79.7% | 3.7% |

| V3-V5 RDP cutoff=80 |  |  |  |  |  |
| --- | --- | --- | --- | --- | --- |
| Rank | AccRDP | OC | MC | Sens | EPQ |
| ----- |  |  |  |  |  |
| p | 99.8% | 0.0% | 0.1% | 99.7% | 0.1% |
| g | 83.3% | 37.8% | 3.4% | 86.0% | 6.6% |

| V3-V5 SINTAX cutoff=90 |  |  |  |  |  |
| --- | --- | --- | --- | --- | --- |
| Rank | AccRDP | OC | MC | Sens | EPQ |
| ----- |  |  |  |  |  |
| p | 99.8% | 0.0% | 0.0% | 99.4% | 0.0% |
| g | 83.2% | 14.7% | 1.3% | 72.5% | 2.6% |

| V3-V5 RDP cutoff=90 |  |  |  |  |  |
| --- | --- | --- | --- | --- | --- |
| Rank | AccRDP | OC | MC | Sens | EPQ |
| ----- |  |  |  |  |  |
| p | 99.8% | 0.0% | 0.1% | 99.6% | 0.1% |
| g | 83.3% | 28.3% | 2.5% | 82.7% | 5.0% |

| V3-V5 SINTAX cutoff=100 |  |  |  |  |  |
| --- | --- | --- | --- | --- | --- |
| Rank | AccRDP | OC | MC | Sens | EPQ |
| ----- |  |  |  |  |  |
| p | 99.8% | 0.0% | 0.0% | 98.0% | 0.0% |
| g | 83.2% | 5.5% | 0.5% | 41.6% | 1.0% |

| V3-V5 RDP cutoff=100 |  |  |  |  |  |
| --- | --- | --- | --- | --- | --- |
| Rank | AccRDP | OC | MC | Sens | EPQ |
| ----- |  |  |  |  |  |
| p | 99.8% | 0.0% | 0.0% | 98.9% | 0.0% |
| g | 83.3% | 13.1% | 1.2% | 67.9% | 2.3% |

| 16S_f1 SINTAX cutoff=50 |  |  |  |  |  |
| --- | --- | --- | --- | --- | --- |
| Rank | AccRDP | OC | MC | Sens | EPQ |
| ----- |  |  |  |  |  |
| p | 99.8% | 75.0% | 0.1% | 99.8% | 0.1% |
| g | 85.6% | 41.6% | 3.2% | 91.7% | 6.8% |

| 16S_f1 RDP cutoff=50 |  |  |  |  |  |
| --- | --- | --- | --- | --- | --- |
| Rank | AccRDP | OC | MC | Sens | EPQ |
| ----- |  |  |  |  |  |
| p | 99.5% | 60.0% | 0.2% | 99.6% | 0.2% |
| g | 85.6% | 78.4% | 4.6% | 94.2% | 11.5% |

16S\_f1 SINTAX cutoff=80
| Rank | AccRDP | OC | MC | Sens | EPQ |
| --- | --- | --- | --- | --- | --- |
| p | 99.8% | 0.0% | 0.0% | 99.7% | 0.0% |
| g | 85.6% | 16.8% | 1.2% | 81.1% | 2.7% |

16S\_f1 RDP cutoff=80
| Rank | AccRDP | OC | MC | Sens | EPQ |
| --- | --- | --- | --- | --- | --- |
| p | 99.5% | 40.0% | 0.1% | 99.5% | 0.2% |
| g | 85.6% | 48.0% | 2.8% | 92.1% | 7.1% |

16S\_f1 SINTAX cutoff=90
| Rank | AccRDP | OC | MC | Sens | EPQ |
| --- | --- | --- | --- | --- | --- |
| p | 99.8% | 0.0% | 0.0% | 99.4% | 0.0% |
| g | 85.6% | 11.2% | 0.9% | 72.7% | 1.8% |

16S\_f1 RDP cutoff=90
| Rank | AccRDP | OC | MC | Sens | EPQ |
| --- | --- | --- | --- | --- | --- |
| p | 99.5% | 26.7% | 0.1% | 99.5% | 0.1% |
| g | 85.6% | 38.6% | 2.3% | 90.8% | 5.7% |

16S\_f1 SINTAX cutoff=100
| Rank | AccRDP | OC | MC | Sens | EPQ |
| --- | --- | --- | --- | --- | --- |
| p | 99.8% | 0.0% | 0.0% | 97.4% | 0.0% |
| g | 85.6% | 3.7% | 0.2% | 39.5% | 0.5% |

16S\_f1 RDP cutoff=100
| Rank | AccRDP | OC | MC | Sens | EPQ |
| --- | --- | --- | --- | --- | --- |
| p | 99.5% | 26.7% | 0.1% | 98.9% | 0.1% |
| g | 85.6% | 21.3% | 1.5% | 84.6% | 3.4% |

ITS\_f1 SINTAX cutoff=50
| Rank | AccRDP | OC | MC | Sens | EPQ |
| --- | --- | --- | --- | --- | --- |
| p | 100.0% | 0.0% | 0.0% | 100.0% | 0.0% |
| s | 75.6% | 54.7% | 8.6% | 85.8% | 15.1% |

ITS\_f1 RDP cutoff=50
| Rank | AccRDP | OC | MC | Sens | EPQ |
| --- | --- | --- | --- | --- | --- |
| p | 99.9% | 100.0% | 0.0% | 100.0% | 0.1% |
| s | 73.9% | 71.6% | 12.5% | 85.0% | 20.8% |

ITS\_f1 SINTAX cutoff=80
| Rank | AccRDP | OC | MC | Sens | EPQ |
| --- | --- | --- | --- | --- | --- |
| p | 100.0% | 0.0% | 0.0% | 99.9% | 0.0% |
| s | 75.6% | 25.6% | 3.2% | 75.2% | 6.4% |

ITS\_f1 RDP cutoff=80
| Rank | AccRDP | OC | MC | Sens | EPQ |
| --- | --- | --- | --- | --- | --- |
| --- | --- | --- | --- | --- | --- |
| p | 99.9% | 0.0% | 0.0% | 99.9% | 0.0% |
| s | 73.9% | 42.2% | 7.9% | 79.8% | 12.7% |

ITS\_f1 SINTAX cutoff=90
| Rank | AccRDP | OC | MC | Sens | EPQ |
| --- | --- | --- | --- | --- | --- |
| --- | --- | --- | --- | --- | --- |
| p | 100.0% | 0.0% | 0.0% | 99.8% | 0.0% |
| s | 75.6% | 18.6% | 2.0% | 69.2% | 4.3% |

ITS\_f1 RDP cutoff=90
| Rank | AccRDP | OC | MC | Sens | EPQ |
| --- | --- | --- | --- | --- | --- |
| --- | --- | --- | --- | --- | --- |
| p | 99.9% | 0.0% | 0.0% | 99.9% | 0.0% |
| s | 73.9% | 31.7% | 6.3% | 76.5% | 9.9% |

ITS\_f1 SINTAX cutoff=100
| Rank | AccRDP | OC | MC | Sens | EPQ |
| --- | --- | --- | --- | --- | --- |
| --- | --- | --- | --- | --- | --- |
| p | 100.0% | 0.0% | 0.0% | 99.2% | 0.0% |
| s | 75.6% | 7.3% | 0.6% | 47.6% | 1.6% |

ITS\_f1 RDP cutoff=100
| Rank | AccRDP | OC | MC | Sens | EPQ |
| --- | --- | --- | --- | --- | --- |
| --- | --- | --- | --- | --- | --- |
| p | 99.9% | 0.0% | 0.0% | 99.6% | 0.0% |
| s | 73.9% | 17.5% | 3.2% | 65.3% | 5.2% |

**Supplementary Table 1.**
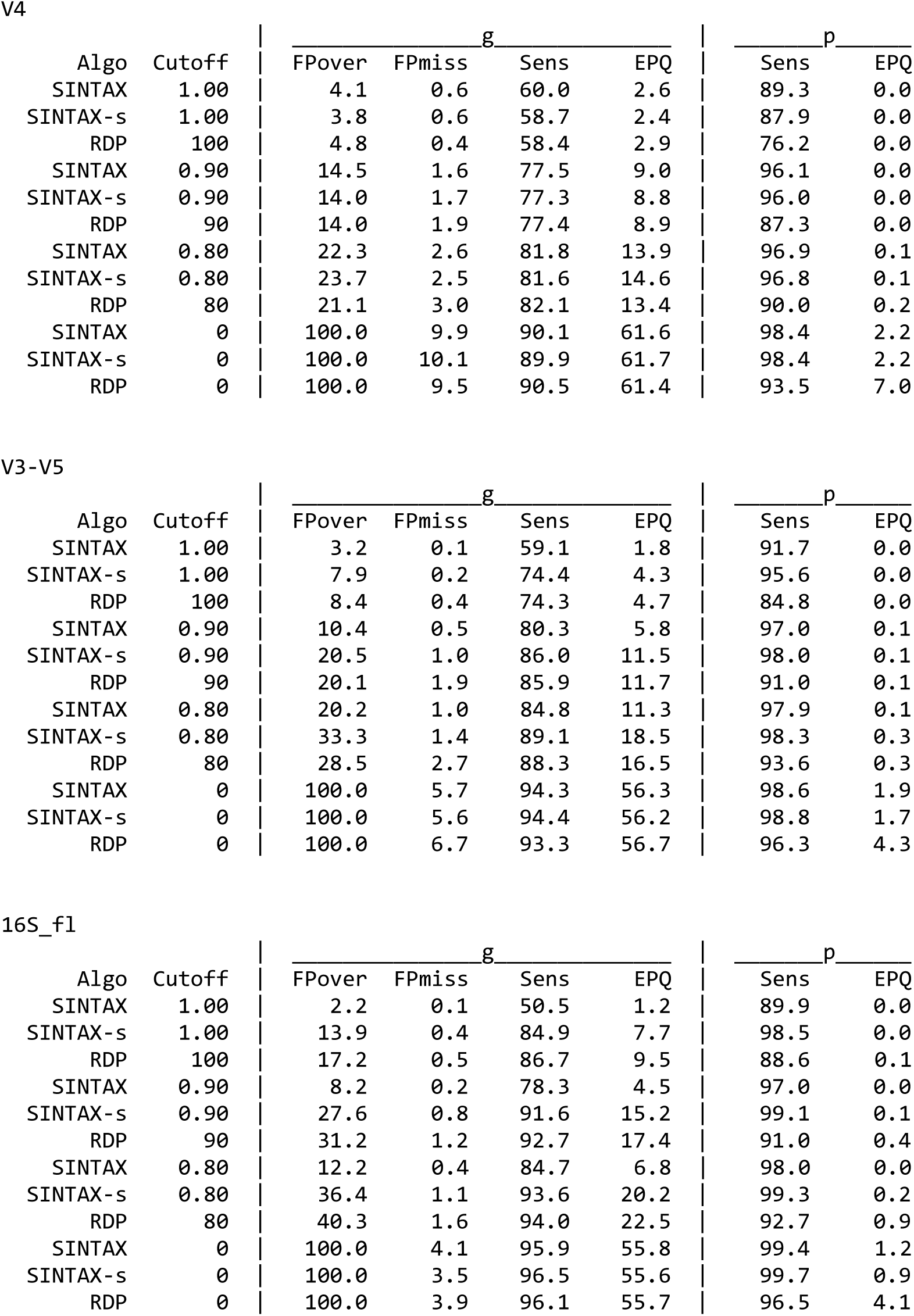
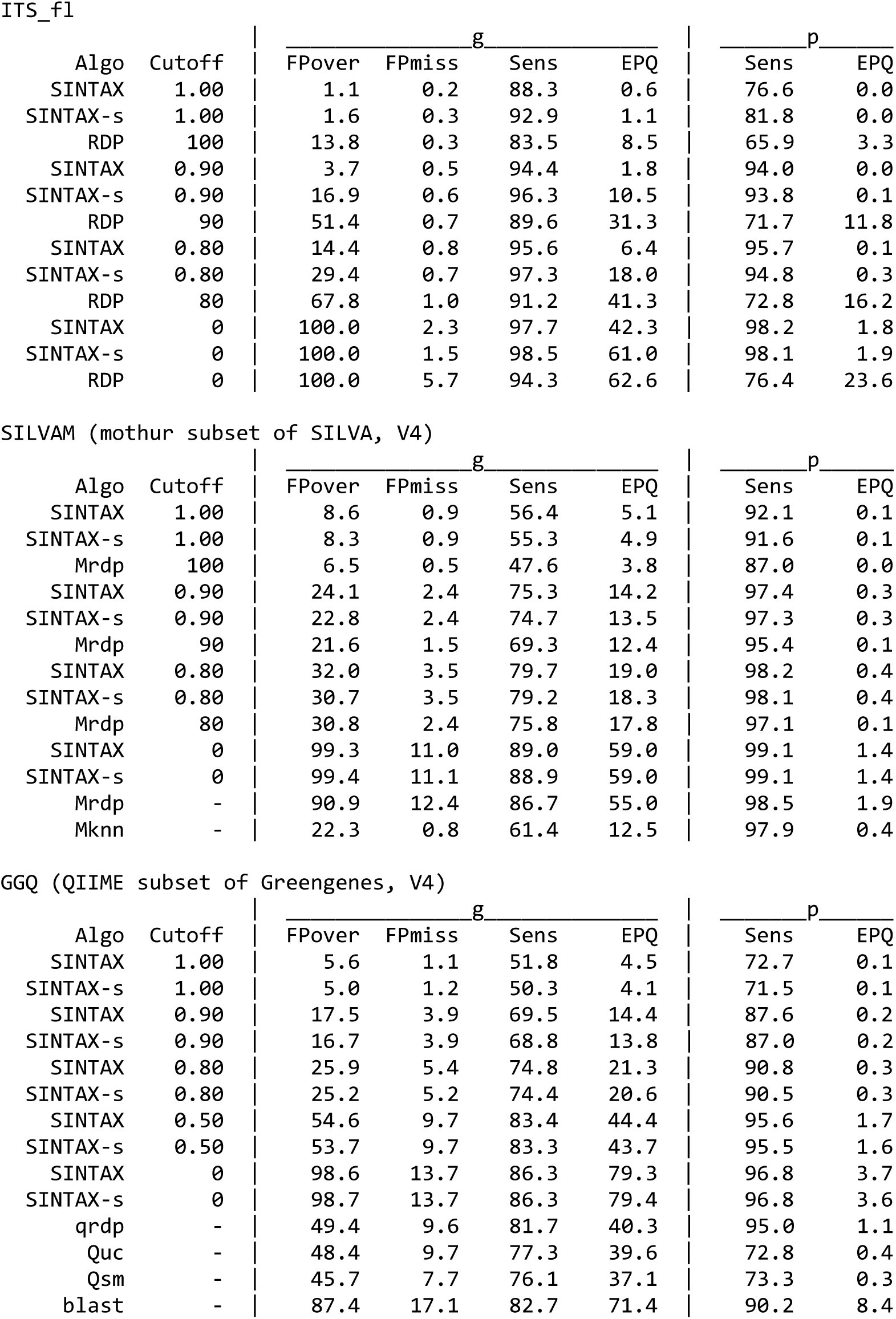
CPX results. SINTAX-s is SINTAX with |Q|/k sub-sample size.

## References

Caporaso, J.G. et al. (2010) QIIME allows analysis of high-throughput community sequencing data. Nat. Methods, 7, 335–336.

DeSantis, T.Z. et al. (2006) Greengenes, a chimera-checked 16S rRNA gene database and workbench compatible with ARB. Appl. Environ. Microbiol., 72, 5069–72.

Deshpande, V. et al. (2015) Fungal identification using a Bayesian classifier and the Warcup training set of internal transcribed spacer sequences. Mycologia, 14–293–.

Edgar, R.C. (2013) UPARSE: highly accurate OTU sequences from microbial amplicon reads. Nat. Methods, 10, 996–8.

HMP Consortium (2012) A framework for human microbiome research. Nature, 486, 215–21.

Kõljalg, U. et al. (2013) Towards a unified paradigm for sequence-based identification of fungi. Mol. Ecol., 22, 5271–5277.

Kozich, J.J. et al. (2013) Development of a dual-index sequencing strategy and curation pipeline for analyzing amplicon sequence data on the miseq illumina sequencing platform. Appl. Environ. Microbiol., 79, 5112–5120.

Lundberg, D.S. et al. (2012) Defining the core Arabidopsis thaliana root microbiome. Nature, 488, 86–90.

Maidak, B.L. et al. (2001) The RDP-II (Ribosomal Database Project). Nucleic Acids Res., 29, 173–4.

McDonald, D. et al. (2012) An improved Greengenes taxonomy with explicit ranks for ecological and evolutionary analyses of bacteria and archaea. ISME J, 6, 610–618.

Pruesse, E. et al. (2007) SILVA: A comprehensive online resource for quality checked and aligned ribosomal RNA sequence data compatible with ARB. Nucleic Acids Res., 35, 7188–7196.

Schloss, P.D. et al. (2009) Introducing mothur: open-source, platform-independent, community-supported software for describing and comparing microbial communities. Appl. Environ. Microbiol., 75, 7537–41.

Schmidt, P.A. et al. (2013) Illumina metabarcoding of a soil fungal community. Soil Biol. Biochem., 65, 128–132.

Wang, Q. et al. (2007) Naive Bayesian classifier for rapid assignment of rRNA sequences into the new bacterial taxonomy. Appl. Environ. Microbiol., 73, 5261–7.

Yarza, P. et al. (2014) Uniting the classification of cultured and uncultured bacteria and archaea using 16S rRNA gene sequences. Nat. Rev. Microbiol., 12, 635–645.

Yilmaz, P. et al. (2014) The SILVA and ‘all-species Living Tree Project (LTP)’ taxonomic frameworks. Nucleic Acids Res., 42.

